# Evaluation of Iron Oxide Nanoparticles for Lymph Node Detection with Magnetomotive Ultrasound – A Pilot Study in Rats

**DOI:** 10.1101/2024.08.28.610049

**Authors:** Ulrika Axelsson, Sania Bäckström, Arefeh Mousavi, Linda Persson

## Abstract

**Introduction:** The inadequate detection of lymph node metastases by current imaging methods has led to overtreatment in rectal cancer. Magnetomotive ultrasound (MMUS) has the potential of being a more accurate diagnostic imaging method for lymph node metastases. This method is based on the detection of tissue movement that is induced by the vibration of iron oxide nanoparticles, caused by an external alternating magnetic field. This study investigated the suitability of the subcutaneous administration of two iron oxide nanoparticles—Ferrotran^®^ and Magtrace^®^—and their distribution in lymph nodes, with regard to the possibility of identifying lymph node metastases in rectal cancer by MMUS.

**Methods:** Male Sprague Dawley rats were injected subcutaneously with 10.7 mg Ferrotran^®^ (n=9), 10.5 mg Magtrace^®^ (n=9), or saline (n=3) dorsally at the root of the tail. On euthanization of the rats after 1, 5, and 24 hours, the proximal and distal lymph nodes were harvested and analyzed by histology [hematoxylin/eosin, Perls Prussian Blue (PPB)] and inductively coupled plasma-optical emission spectroscopy. The primary aim was to evaluate the distribution of the iron oxide nanoparticles throughout the lymphatic system; the general health of the rats after subcutaneous administration of these nanoparticles was also monitored.

**Results:** At 1 hour after subcutaneous injection, Ferrotran^®^ and Magtrace^®^ accumulated in the proximal lymph nodes. After 24 hours, both particles had spread to distal lymph nodes, but only Ferrotran^®^ reached the mesenteric and mandibular lymph nodes. In addition, Ferrotran^®^ penetrated the lymph nodes more deeply than Magtrace^®^ at 24 hours. No toxicity was observed with either nanoparticle.

**Conclusion:** Although both compounds disseminated well, Ferrotran^®^ accumulated better and more rapidly in lymph nodes than Magtrace^®^. Because accumulation and time are important parameters for imaging, our data indicate that Ferrotran^®^ is a potentially more suitable particle for MMUS in clinical use.

## Introduction

Colorectal cancer is one of the most common cancers, with 732,000 new cases worldwide in 2020 [1]. Nevertheless, survival in colorectal cancer has increased in recent decades due to improved treatment, changing patterns in risk factors, and better screening, with 5-year survival rates reaching 67% [2,3]. The presence of lymph node metastases is a key determinant of the prognosis of rectal cancer (and cancers in general), guiding the development of an adequate treatment plan. Combined with computed tomography (CT), biopsy, and colonoscopy/rectoscopy, lymph node status is generally assessed by magnetic resonance imaging (MRI) [4], F-18 fluorodeoxyglucose positron emission tomography (PET)/CT, or PET/MRI [5,6]. Finally, lymphatic flow and regional lymph nodes can be visualized by radionuclide scintigraphy in various cancers.

These methods, however, are associated with several drawbacks. MRI is insufficient for diagnosing lymph node status, because it only analyzes the shape and size of lymph nodes. Large lymph nodes could be suspected of metastases and treated as such, but they might result from inflammation or constitute bigger nodes that are benign. Conversely, smaller lymph nodes will not appear to be questionable, based on size, and will thus be overlooked. Consequently, the diagnostic accuracy of these methods for small metastatic lymph nodes (< 5 mm) declines substantially. In addition, scintigraphy is suitable for identifying sentinel lymph nodes but can not detect the presence or absence of cancer in them [7]. In contrast, PET can differentiate cancerous versus noncancerous lymph nodes [8] but is expensive to perform. Thus, current methods detect lymph node metastases suboptimally or have limited widespread use, necessitating a more tenable approach to identifying lymph nodes for diagnostic purposes.

Further, due to the unavailability of a suitable technique, it is challenging to determine whether a tumor has spread to regional lymph nodes, forcing most patients to undergo total mesorectal excision (TME) by default. However, studies have shown that nearly 90% of patients with stage T1 who underwent TME experienced no spreading of the tumor to lymph nodes and thus would not have needed this extensive surgery [9]. This overtreatment negatively impacts quality of life and raises healthcare costs, necessitating more effective tools to determine lymph node status in rectal cancer and address this significant unmet need.

Magnetomotive ultrasound (MMUS) is a novel diagnostic imaging method that has tremendous potential as a platform for making more accurate diagnoses [10]. MMUS is based on the detection of vibrating iron oxide nanoparticles in tissue. Through application of a time-varying magnetic field, the nanoparticles and thus the immediate surrounding tissue are set in motion [11,12]. Theoretically, in this modality, conventional US is first performed to generate a US image. When an alternating magnetic field is applied, the nanoparticles within the tissue begin to vibrate and set the tissue in motion; this activity is detected by US, yielding a contrast image at a resolution of millimeters. Because this motion is detected by US, the low echogenicity of nanocompounds can be overcome [13], generating better images and improving the diagnostic accuracy. In addition, nanoparticles are transported primarily by phagocytosing macrophages; these macrophages do not enter tumor cells. Thus, the tumor cells in a lymph node will harbor limited amounts of nanoparticles, resulting in less vibration—a difference that can be distinguished by MMUS, differentiating between lymph nodes with and without tumor cells, beyond the imaging of sentinel lymph nodes that is accomplished by other techniques. Whether MMUS can make this differentiation in a clinical setting remains to be confirmed in clinical studies.

Further, MMUS devices have been developed to be smaller and cheaper and to require less training than MRI, with the goal of recording smaller lymph nodes and metastases down to 2 mm.

To detect lymph node metastases by MMUS, iron oxide nanoparticles should be injected into the tissue and spread to the lymph nodes. Notably, the nanoparticles should have sufficient time to reach the lymph nodes before the analysis is performed. Further, the nanoparticles should not be toxic at the administered dose.

Two superparamagnetic iron oxide nanoparticles (SPIOs) are now being examined with regard to their suitability for MMUS in clinical practice. Ferrotran^®^, an ultrasmall superparamagnetic iron oxide (USPIO) particle, is an effective intravenously administered contrast agent for detecting lymph node metastases by MRI in various cancers [14–17]. Ferrotran^®^ is currently in a Phase III study for improving MRI images in prostate cancer. Magtrace^®^ is an SPIO (Endomag, Cambridge, UK) that has been approved by the FDA as a sentinel lymph node tracer in breast cancer patients.

The distribution of intravenous injection of Ferrotran^®^ has been studied using MRI for detecting lymph node metastases in various cancers, including rectal cancer [16,18,19]. The assessment of lymph nodes in rectal cancer by MMUS entails local injection of the tracer inside the rectal lining of the intestine, around the tumor. Consequently, subcutaneous delivery of nanoparticles, which will result in a different biodistribution than intravenous injection—thus necessitating distinct examination windows and doses—has potential application in diagnosing rectal cancer stage using MMUS. Although smaller nanoparticles might be expected to distribute more efficiently within the lymphatic system, there is no evidence that nanoparticle size is the sole determinant of the rate of distribution. Further, given the disparate structures between Ferrotran^®^ and Magtrace^®^, it would be premature to assume that the former would circulate to a greater extent, prompting us to determine which of the latter disseminates better for use in MMUS.

The aim of this study was to examine the spread and distribution of two iron oxide nanoparticles— Ferrotran^®^ and Magtrace^®^—to lymph nodes in biological tissue, after subcutaneous administration in male Sprague Dawley rats. This model was chosen, based on size considerations: their lymph nodes are approximately the same size as human mesorectal lymph nodes, and the rat is roughly the same length as the mesorectum [16], ensuring that we are studying a relevant distance from the injection site to the lymph nodes. As a secondary aim, we monitored the general health of the rats after local subcutaneous administration of Ferrotran^®^ and Magtrace^®^ to confirm the reported safety of these nanoparticles. Thus, the overarching purpose of this study was to obtain an initial indication of which tracer is more suitable for diagnostic imaging with MMUS.

## Materials and methods

### Animals

Male Sprague Dawley rats were obtained from Janvier Labs (France). At the time that the doses were administered, the rats weighed approximately 200–250 g. The rats were housed in type III European standard cages (3 rats per cage) with Aspen wood chip bedding (Tapvei, Brogaarden, Denmark). Each cage contained a hide and chewing stick for environmental enrichment. The temperature was held between 20 °C and 24 °C, and the humidity was set to 50% to 65% RH.

On the day of arrival, the animals were picked randomly from the crates and allocated successively to the test groups. No randomized allocation sequence was used. Before administration of the test compounds, all animals underwent an acclimatization period during which they were observed for signs of ill health; any animal in poor condition was excluded. Ultimately, no animals were excluded from the study. After administration of the test compounds, all animals were inspected throughout the study with regard to their general condition. All clinical signs and behavioral abnormalities were recorded.

The investigators were not blinded to the group allocation during the experiment, outcome assessment, or data analysis. This study was conducted in accordance with license numbers 18164-21 and M140-16 per the Malmö/Lund Ethics Committee on Animal Testing.

### Test compounds

This study assessed the iron oxide nanoparticle solutions Magtrace^®^ (Endomag, Cambridge, UK) and Ferrotran^®^ (ferumoxtran; SPL Medical, Nijmegen, Netherlands). Ferrotran^®^ (ferumoxtran) is being developed by SPL Medical as an MRI contrast agent for detecting lymph node metastases. Ferrotran^®^ comprises 25-nm iron oxide nanoparticles with a 4-nm-diameter core of Fe_3_O_4_ and a dextran coating and is stabilized with sodium citrate. Magtrace^®^, developed by Endomag, is used for breast cancer staging by assisting in local lymph node detection at the tumor site, as part of a sentinel lymph node biopsy. Magtrace^®^ is composed of 60-nm iron oxide nanoparticles with a 4-nm-diameter core of Fe_2_O_3_ and Fe_3_O_4_ and a carboxydextran coating and is stabilized with sodium chloride.

Two batches of Ferrotran^®^—batch 1 (B3008435) and batch 2 (B3008435)—consisting of a lyophilized powder (0.5 g of powder), were reconstituted in vials with 5 ml 0.9% NaCl solution to a concentration of 19.9 mg/ml. The vials were inverted several times to ensure that a homogeneous solution was obtained (no vortexing). After reconstitution, the solutions of iron oxide nanoparticles were uniformly dark brown to black. The 2 batches were stored at room temperature until reconstitution and injection. All injections of the reconstituted test compound were performed on the same day.

One batch of Magtrace^®^ (1114Ph112), supplied as a solution (28 mg Fe/ml), was diluted in 0.9% NaCl. To ensure that a homogeneous solution was obtained, the vial was inverted several times (no vortexing) to a concentration of 21.2 mg Fe/ml. Prior to filling the syringes, all solutions were mixed by gentle swirling to ensure homogeneity.

### Dosing

Sprague Dawley rats were administered subcutaneously, dorsally at the root of the tail, with Ferrotran^®^, Magtrace^®^, or saline (9 mg/ml) (stored at 4 °C until use; Timeline Bioresearch AB) (Table 1). No formal a priori sample size calculation was performed to select the number of rats per treatment group, because this qualitative exploratory study was intended to provide initial insights into the spread of the test compounds.

**Table 1.**
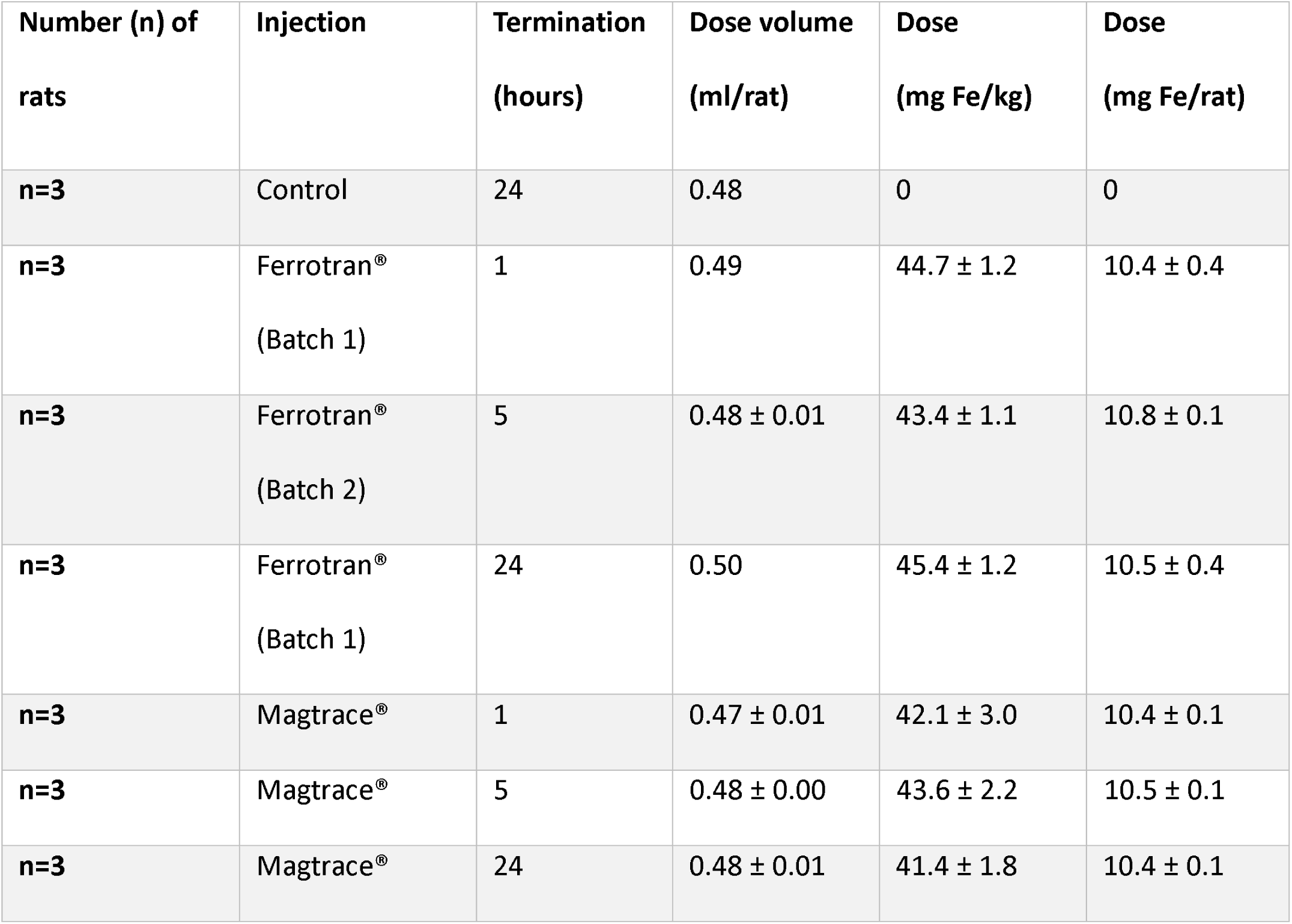
Doses of test compounds.

| Number (n) of rats | Injection | Termination (hours) | Dose volume (ml/rat) | Dose (mg Fe/kg) | Dose (mg Fe/rat) |
| --- | --- | --- | --- | --- | --- |
| n=3 | Control | 24 | 0.48 | 0 | 0 |
| n=3 | Ferrotran®<br>(Batch 1) | 1 | 0.49 | 44.7 ± 1.2 | 10.4 ± 0.4 |
| n=3 | Ferrotran®<br>(Batch 2) | 5 | 0.48 ± 0.01 | 43.4 ± 1.1 | 10.8 ± 0.1 |
| n=3 | Ferrotran®<br>(Batch 1) | 24 | 0.50 | 45.4 ± 1.2 | 10.5 ± 0.4 |
| n=3 | Magtrace® | 1 | 0.47 ± 0.01 | 42.1 ± 3.0 | 10.4 ± 0.1 |
| n=3 | Magtrace® | 5 | 0.48 ± 0.00 | 43.6 ± 2.2 | 10.5 ± 0.1 |
| n=3 | Magtrace® | 24 | 0.48 ± 0.01 | 41.4 ± 1.8 | 10.4 ± 0.1 |

The rats were administered the test compounds under isoflurane anesthesia. The injection site was shaved before injection. To determine whether increased blood and lymphatic circulation in the area affects the spread of the particles, the injection site was massaged gently for roughly 2 min approximately 90 min after the injection for 1 rat per treatment and time point (6 rats in total). The actual dose volumes were determined by weighing the syringes prior to and after administration.

Termination Rats were terminated by CO_2_ inhalation per the schedule in Table 1. After termination, the rats were shaved and photographed using a mobile phone camera. The diameter of the area of discolored skin around the injection site was measured.

### Tissue collection and handling

The rats were opened and dissected 1–6 hours after termination to expose the lymph nodes. The appearance of the organs, lymph nodes, and lymphatic vessels was noted. The following lymph nodes were taken from both sides when possible: superficial inguinal, external iliac, deep inguinal, mesenteric, axillary, and mandibular. Lymph nodes and the tissue in and around the injection site were dissected with a scalpel, photographed, and analyzed per Table 2. In some cases, 1 lymph node (left or right) was processed for inductively coupled plasma-optical emission spectrometry (ICP-OES), and the other node was fixed for histology. Lymph nodes for ICP-OES were weighed, frozen at -20 1C, and stored at -80 1C, and those for histology were fixed in 4% buffered formalin (see Histology). Some lymph nodes could not be collected for technical reasons and are therefore missing.

**Table 2.** Sample handling.

|  |  | Number of rats sampled per analysis |  |  |
| --- | --- | --- | --- | --- |
| Injection | Termination<br>(hours) | Lymph<br>nodes for<br>ICP-OES | Lymph nodes<br>for histology | Injection site<br>tissue sample<br>for histology |
| Control | 24 | n=2 | n=2 | n=0 |
| Ferrotran® (Batch 1) | 1 | n=0 | n=3 | n=1 |
| Ferrotran® (Batch 2) | 5 | n=2 | n=2 | n=0 |
| Ferrotran® (Batch 1) | 24 | n=0 | n=3 | n=1 |
| Magtrace® | 1 | n=2 | n=3 | n=1 |
| Magtrace® | 5 | n=2 | n=2 | n=0 |
| Magtrace® | 24 | n=2 | n=2 | n=1 |

### Histology

Tissues were delivered to Micromorph AB (Lund, Sweden) in 4% buffered formalin. The samples were dehydrated, cleared, infiltrated with paraffin in an automated TISSUE-TEK V.I.P. (Miles Scientific, Newark, US), and embedded in paraffin. Sections (4 μm) were prepared and dried in an oven at 37 °C overnight.

#### Hematoxylin and Eosin

Sections were deparaffinized and hydrated in distilled water, after which they were stained in Mayeŕs hematoxylin (BioOptica, Milano, Italy, 05-6002) for 6 minutes, washed in running tap water for 10 minutes, and washed in distilled water for several seconds. Next, they were stained in eosin (BioOptica, 05-11007) for 3 minutes, washed in distilled water for several seconds, and dehydrated sequentially in 95% EtOH and 100% EtOH. The sections were then cleared and mounted in synthetic resin.

#### Iron Staining-Perls Prussian Blue

The Perls Prussian Blue kit (ab150674, Abcam, Cambridge, UK) was used to stain iron. Tissue sections were deparaffinized and hydrated in distilled water. Equal volumes of potassium ferrocyanide solution and hydrochloric acid solution were mixed to establish a working iron stain solution. Slides were incubated in iron stain solution for 3 minutes and rinsed thoroughly in distilled water. The slides were then stained in Nuclear Fast Red Solution (Abcam) for 5 minutes and rinsed in 4 changes of distilled water. Finally, they were dehydrated sequentially in 95% EtOH and 100% EtOH, cleared, and mounted in synthetic resin.

#### Analysis

The sections were analyzed and photographed under a Leica DMRX microscope. An experienced assessor estimated the level of iron staining, expressed as the percentage of lymph node area and lymph node circumference that was occupied by iron-positive cells. Hematoxylin/eosin-stained sections were used as a reference for the PPB iron staining.

### ICP-OES bioanalysis

Briefly, after being weighed, the tissues were stored at -80 °C. The tissue was thawed and digested in Teflon vials (Prototyp verkstaden, Medicon Village, Lund) with 38.08% nitric acid, HNO_3_ (Normatom, VWR Chemicals), 13.2% hydrogen peroxide, H_2_O_2_ (Sigma Aldrich, Missouri, USA), and Milli-Q water (Milli-Q Plus 185, resistivity: 18.2 MΩ x cm). The samples were heated for 90 minutes at 100 °C with shaking at 240 rpm and cooled for at least 30 min. Two milliliters of each sample was then diluted to 5 ml 15.2% HNO_3_ and 5.3% H_2_O_2_ before analysis.

The ICP-OES analysis was performed on an ICP-OES 710 that was equipped with a quartz torch with a 2.4-mm injector, a single-pass glass cyclonic spray chamber, and a OneNeb inert concentric or glass nebulizer (all from Agilent, Santa Clara, USA). The instrument settings were as follows: number of replicates, 3; sample uptake delay time, 40 s; stabilization delay, 20 s; replicate read time, 5 s; plasma gas flow, 15 L/min; auxiliary gas flow, 1.5 L/min; RF power, 1.2 kW; nebulizer flow, 0.75 L/min; and flow rate of peristaltic pump, 15 rpm. A weighted linear calibration curve was generated, with a maximum 5% error allowed.

### Statistics

Results are presented as representative images that show the relative amounts and locations of the nanoparticles. Values from semiquantitative analyses are presented using descriptive statistics—ie, the mean and standard deviation—and graphed as bar plots. No significance testing was applied due to the qualitative nature of this study.

## Results

### Toxicology

The toxicity of the injected nanoparticles to the general health of all rats was followed until termination (see Table 1), for a maximum of 24 hours. No clinical or behavioral abnormalities were noted in any rat during this time, and all organs appeared normal.

### Spread of nanoparticles from site of injection

Sprague Dawley rats were injected with Ferrotran^®^ or Magtrace^®^ solution (10.3–10.9 mg Fe) or saline dorsally at the root of the tail. The injection site turned brown-black after the administration of Ferrotran^®^ and Magtrace^®^ (Figure 1), as expected due to the iron—a change that was not observed in the control group. The discoloration of the injection site did not increase in size from 1 to 24 hours post-dose with Magtrace^®^. However, with Ferrotran^®^, the brown area spread visibly from the injection site, increasing in diameter from 10 mm 1 hour after injection to 25 mm 24 hours post-dose.

**Figure 1.**
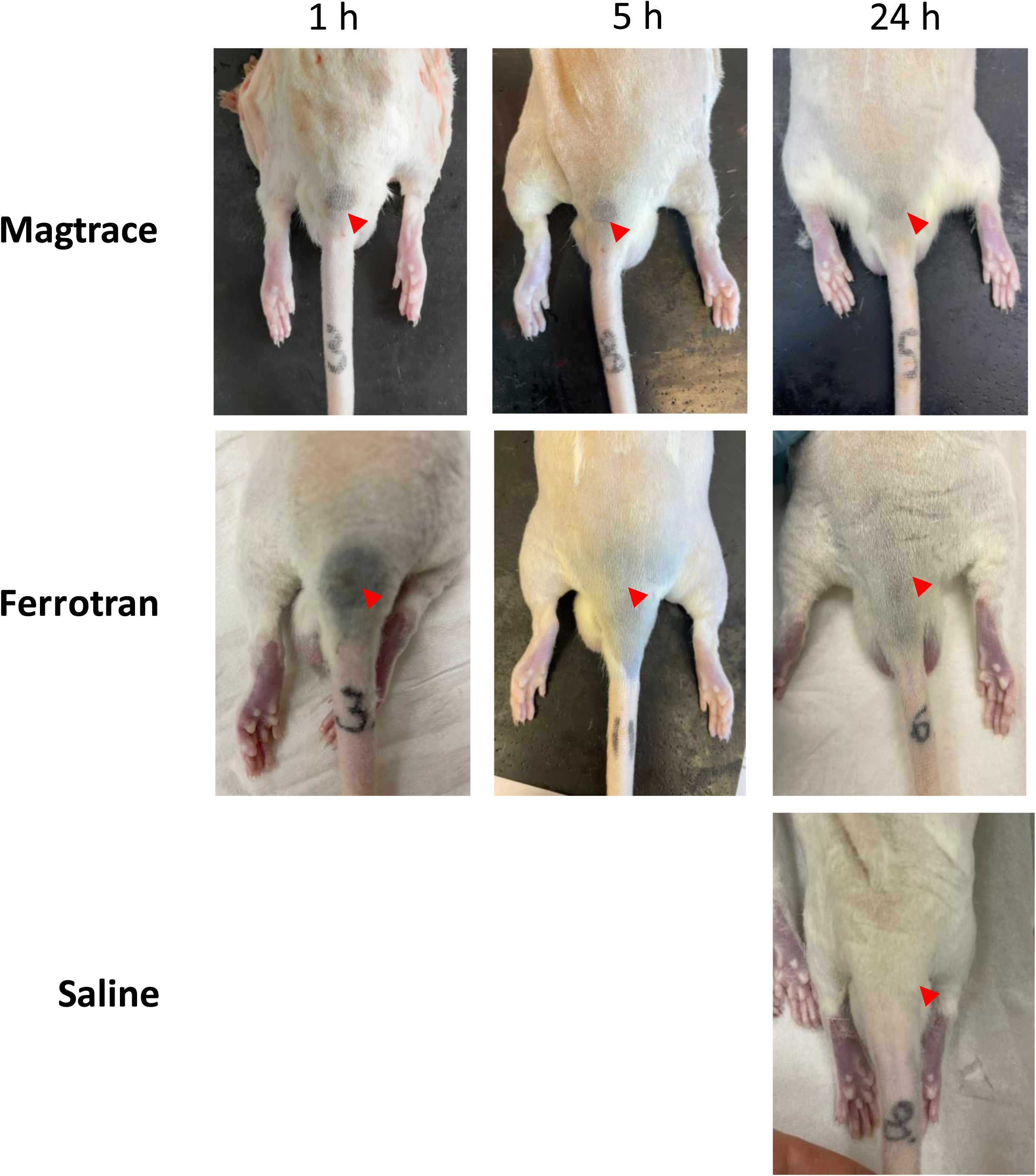
Representative photographs of injection sites at 1, 5, and 24 hours after administration of Magtrace^®^ and Ferrotran^®^. The red arrows denote the injection site.

### Spread of nanoparticles to lymph nodes

Lymph nodes can be grouped as proximal and distal from the injection site. In rat, we defined the superficial inguinal, deep inguinal, and external iliac lymph nodes as proximal and the mesenteric, axillary, and mandibular lymph nodes as distal (Figure 2).

**Figure 2.**
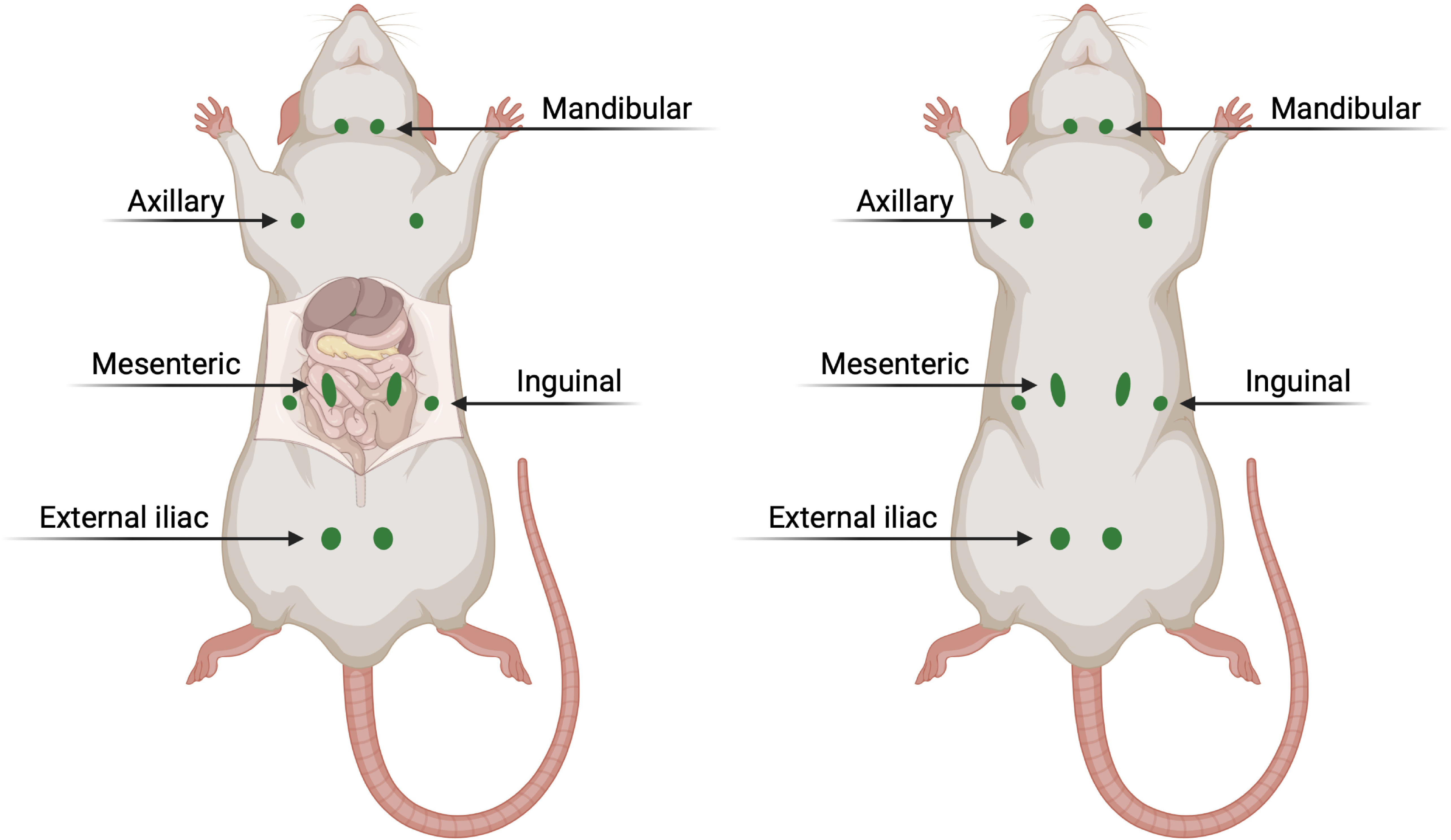
Location of proximal and distal lymph nodes in rat. Created with BioRender.com.

The proximal lymph nodes contained nanoparticles 1 hour after injection, visible by eye (Figure 3A). The nanoparticles also resided in lymphatic vessels, as evidenced by darkening of the lymphatic vessels between the superficial inguinal and axillary nodes, indicating that these tracers migrate between nodes through the lymphatic vessels (Figure 3C). Further, the lymph nodes darkened after 24 hours versus 1 hour post-dose, indicating that nanoparticles accumulated in the lymph nodes after injection. After 24 hours post-dose, the nanoparticles reached the axillary lymph nodes. Ferrotran^®^ also entered the mesenteric lymph nodes by gross photography and histology (Figures 3B and 5B, respectively); entry into the mandibular lymph nodes was evident only by histology (Figure 5B). A more homogeneous distribution of nanoparticles was seen in the lymph nodes near the injection site after 24 hours compared with 1 hour.

**Figure 3.**
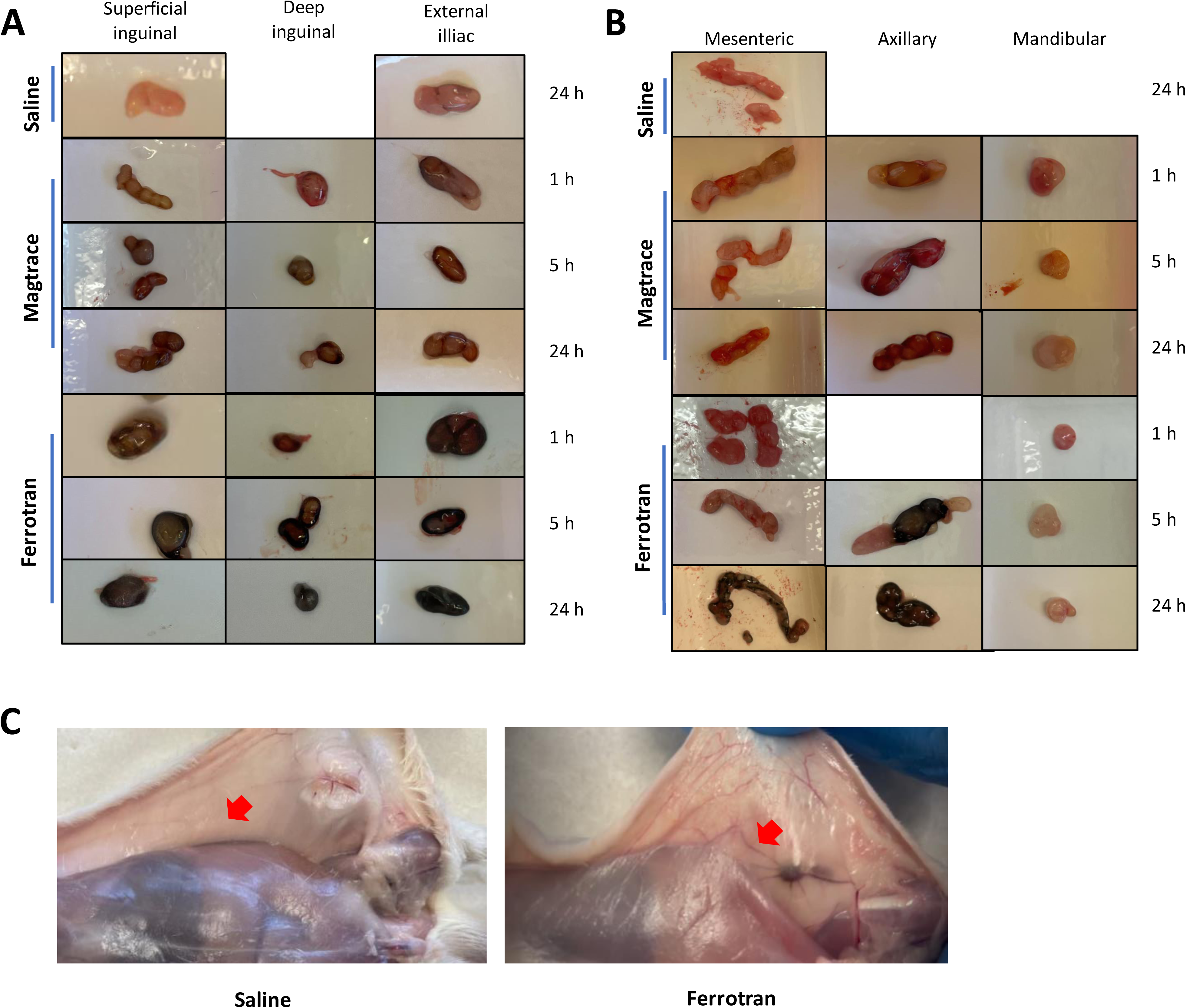
Representative photographs of A. proximal and B. distal lymph nodes in rats 1, 5, and 24 hours after administration of Ferrotran^®^ and Magtrace^®^ versus saline control. Some lymph nodes could not be collected for technical reasons and are therefore missing. C. Representative images of lymphatic vessels in rats 24 hours after injection with Ferrotran^®^ versus saline control. As indicated by the arrows, the lymph vessels are stained for iron by Ferrotran^®^ nanoparticles.

### Distribution of nanoparticles in lymph nodes

To evaluate their distribution within lymph nodes and the injection site, the nanoparticles were stained with PPB. There was no nonspecific staining of iron in the lymph nodes or injection site in control rats. However, at the injection site in the animals that were injected with nanoparticles, the nanoparticles spread to cells and fibrous structures in the dermis after injection, as shown in Figure 4A-B.

**Figure 4.**
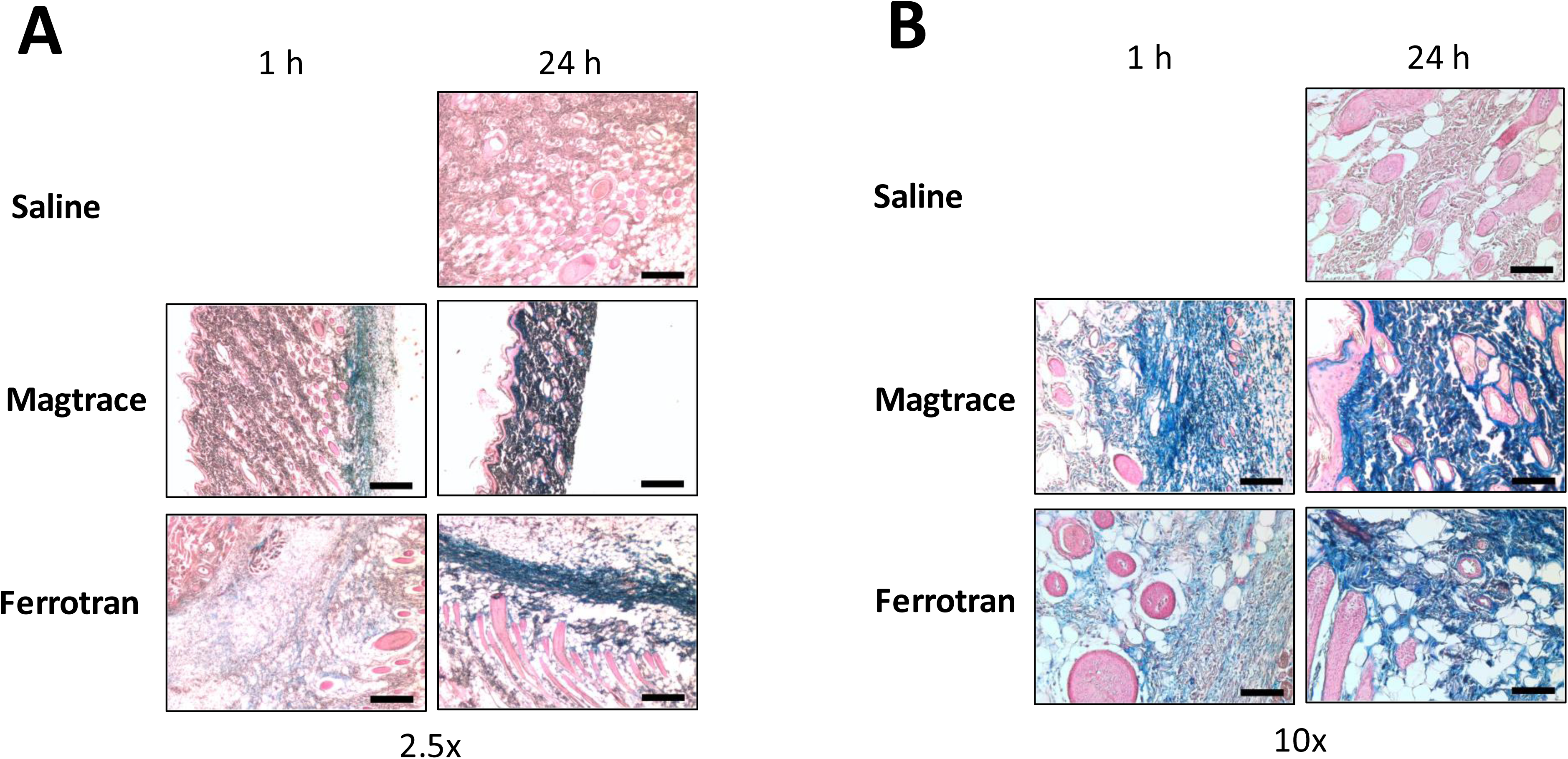
Histological analysis of skin and dermis at the injection site 1 and 24 hours after administration of Ferrotran^®^ and Magtrace^®^ by Perls Prussian Blue staining. A. 2.5x and B. 10x magnification. The blue cells indicate the presence of iron. Scale bar in A. and B. equals 100 μm and 400 μm, respectively.

Representative PPB-stained images of proximal and distal lymph nodes are shown in Figure 5A and 5B, respectively, and the estimated percentages of the lymph node area and circumference that were stained by PPB are shown in Figure 6A and 6B, respectively. Based on the stains, iron accumulation increased in proximal lymph nodes from 1 to 5 to 24 hours. At 24 hours post-dose, Ferrotran^®^ filled 10% to 55% of the lymph node area (except for the mandibular and mesenteric lymph nodes), versus 2% to 30% with Magtrace^®^ (Figure 6A). The mandibular lymph nodes, which can only be reached by nanoparticles through the blood, did not contain particles at 1 hour and 5 hours post-dose but appears to have harbored Ferrotran^®^ particles at 24 hours post-dose; however, no accumulation of Magtrace^®^ was observed. Unlike the weak histological and photographic evidence of the entry of Ferrotran^®^ into the mesenteric lymph nodes in Figures 3B and 5B, no such distribution was observed by PPB staining (Figure 6A and B); the low coverage of PPB staining in the mandibular lymph nodes by Ferrotran^®^ is consistent with the weak staining by histology (Figure 6B vs Figure 5B). Overall, these data demonstrate the potential uptake and transportation of Ferrotran^®^ by the blood in rat within 24 hours (Figure 5B).

**Figure 5.**
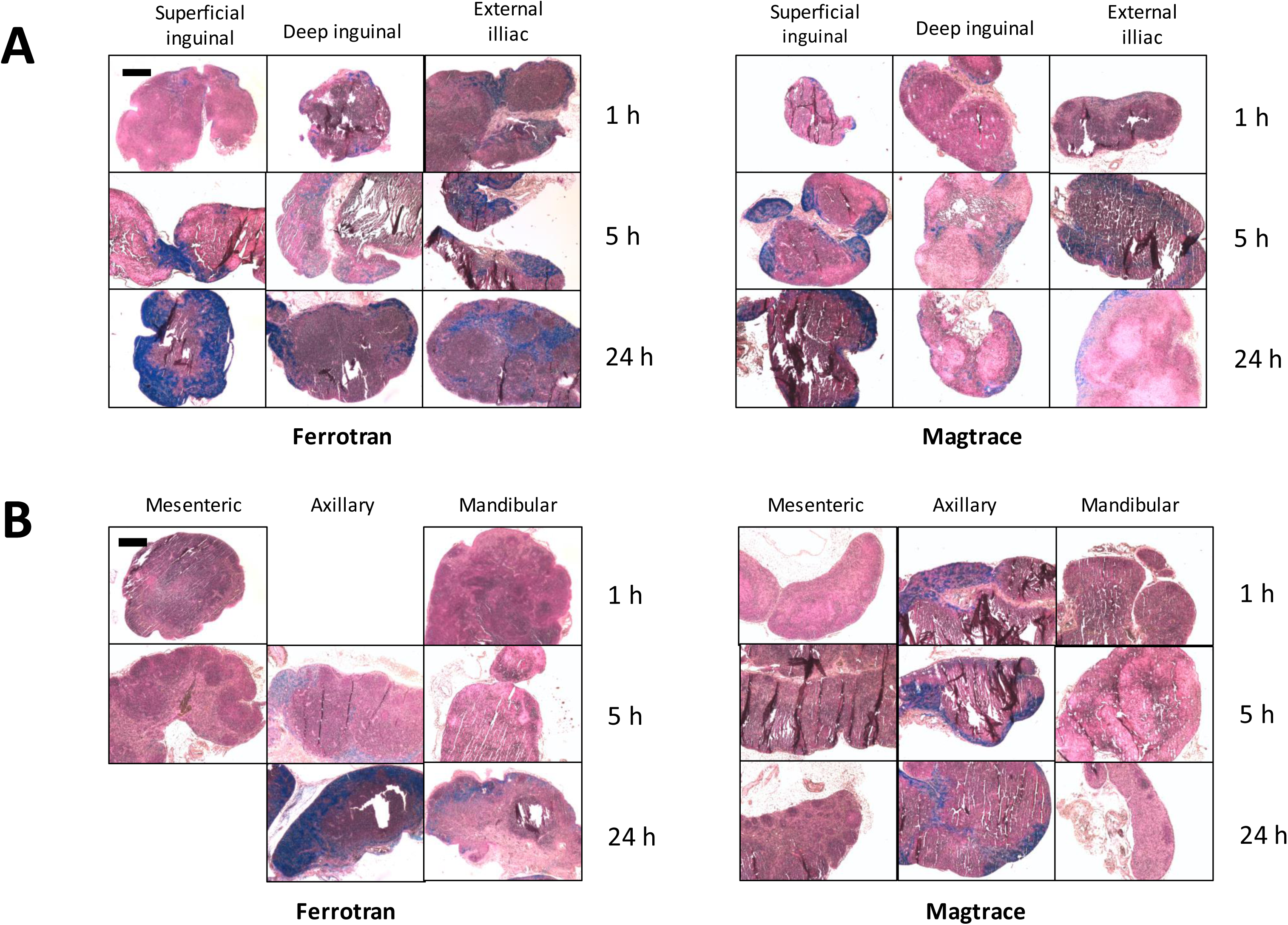
Histological images of Perls Prussian Blue-stained A. proximal and B. distal lymph nodes after administration of Ferrotran^®^ and Magtrace^®^. Blue signals indicate the presence of nanoparticles. Some lymph nodes could not be collected for technical reasons and are therefore missing. Scale bars (black lines in top left panels) equal 400 μm.

**Figure 6.**
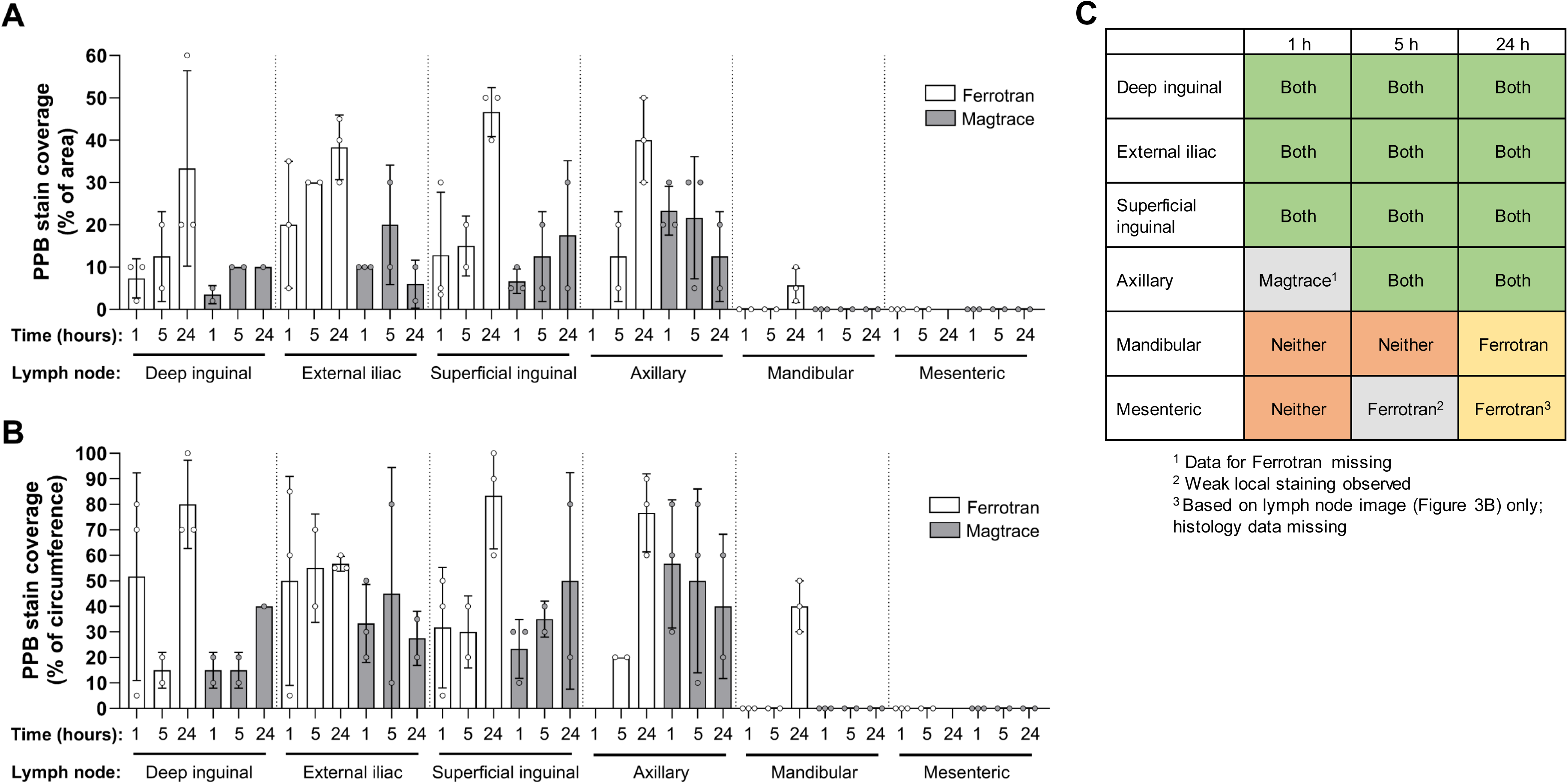
Estimated percentage of A. lymph node area and B. lymph node circumference stained by Perls Prussian Blue. Bars are the mean, and whiskers are the standard deviation. For rats that had 2 of the same type of lymph node collected, the nodes were considered technical replicates, and their mean was plotted. Some lymph nodes could not be collected for technical reasons and are therefore missing. C. Summary of the presence or absence of each tracer in the various lymph nodes at each time point.

Further, as shown by the PPB stain at 1 hour post-dose, the nanoparticles reached the subcapsular sinus (ie, the periphery of the lymph node), indicating rapid transport to proximal lymph nodes. However, after 24 h, the nanoparticles penetrated deeper into lymph nodes and spread more homogeneously within them compared with 1 h; this pattern was more evident for Ferrotran^®^ than for Magtrace^®^. A summary of the lymph nodes that were reached by each nanoparticle at each time point is presented in Figure 6C.

The concentration of iron in the lymph nodes was measured by ICP-OES. Unfortunately, we had too few lymph nodes to analyze to generate any significant calculations. However, generally, the concentrations of iron in lymph nodes were higher over time after injection. After 5 hours (the only time point that was available for both nanoparticles), the concentration of iron was higher in lymph nodes in rats that were injected with Ferrotran^®^ versus Magtrace^®^, indicating that Ferrotran^®^ spreads faster and more broadly; however, we had too little data to draw any conclusions.

### Transport mechanism

In the lymph nodes, iron was found primarily in macrophage-like cells but also between cells, possibly in the lymph node sinuses. These findings suggest that nanoparticles are transported primarily by macrophages, as reported [20]. In addition, the macrophage-like cells that contained iron were larger at 24 hours versus 1 hour post-dose (Figure 7), indicating continuous uptake of the tracer by these cells.

**Figure 7.**
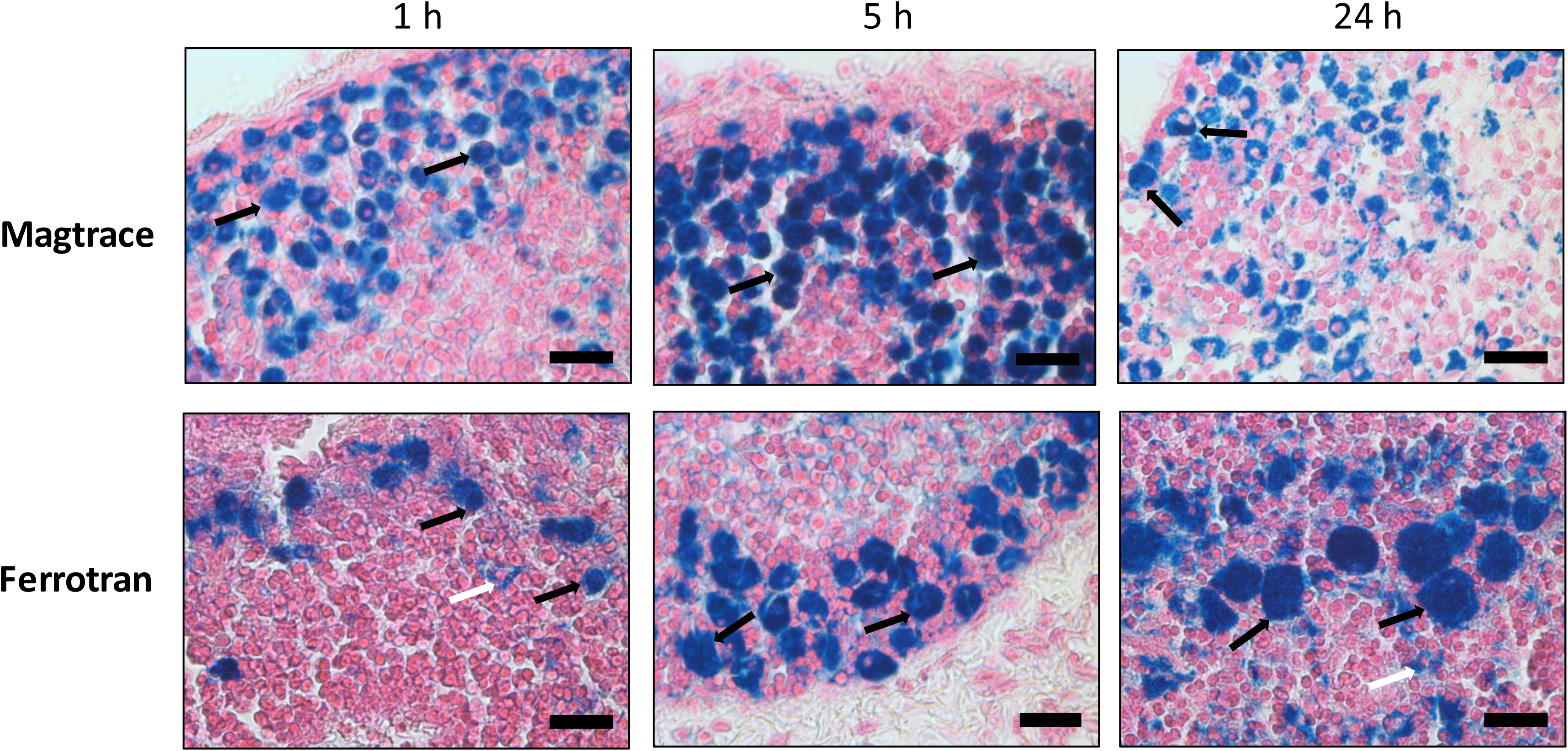
Increase in size of macrophage-like cells over time after administration of Ferrotran^®^ by PPB staining. At 24 h, diffuse staining is also observed between cells. Scale bar equals 20 μm.

### Lymph node size

The lymph nodes in rat (mean 3.2 mm; range 1.2–6.6 mm) approximate the mesorectal lymph nodes in human in size (mean 2.4 mm; range 0.9–12.2 mm) [16]. In our study, the long axis of lymph nodes in the rats ranged from 1.4 to 7.3 mm, and the short axis ranged from 1 to 3.3 mm.

### Influence of massage on nanoparticle spread

There was no obvious difference in the spreading of nanoparticles between rats that had been massaged or not—nanoparticles entered the lymphatic system and spread to distal lymph nodes even without massage.

## Discussion

In this study, we have found that on subcutaneous injection, Ferrotran^®^ and Magtrace^®^ iron oxide nanoparticles enter the lymphatic system and are transported to proximal and distal draining lymph nodes in rat. Although both compounds disseminated well, Ferrotran^®^ accumulated better and more rapidly in lymph nodes than Magtrace^®^. Neither nanoparticle was associated with any gross clinical or behavioral toxicity.

In rectal cancer, the status of regional and mesorectal lymph nodes is central for staging the cancer and predicting local and distant recurrence—this information will inform treatment decisions with regard to preoperative diagnostic imaging, surgical techniques, pathological assessment, and the use of radiation therapy. Although sentinel lymph nodes have been analyzed in rectal cancer patients, there is no consensus on whether they exist [21,22]. Despite some studies disputing their presence, others have reported sentinel lymph nodes to lie 1–10 cm from the primary tumor in colorectal cancer patients [23]. Thus, one must also be able to analyze lymph nodes that are distal to the tumor.

MMUS is a novel, potentially suitable technique that entails US under the application of an external magnetic field. An MMUS system locates lymph nodes by high-resolution US and tracks the distribution of a contrast agent throughout lymph nodes [10,24]. MMUS has the potential to provide images with higher resolution compared with MRI in clinical settings and has the significant benefit of being free of ionizing radiation. In addition, lymph nodes in the rectal area, which are often difficult to evaluate due to anatomical circumstances, can be examined via the rectum. Thus, a small, easy-to-use, portable MMUS-based rectal US probe would potentially have a great impact in diagnosing rectal cancer, benefiting patients.

Iron-oxide nanoparticles have been used in various studies for detecting lymph nodes and lymph node metastases [14–17,19] but have only recently been applied in MMUS. Jansson *et al*. reported the first MMUS image of human tissue using iron oxide nanoparticles in 2023, in which a patient was injected with nanoparticles prior to surgery, and an excised tissue specimen was imaged on a prototype MMUS system (NanoEcho, Lund, Sweden) [13].

Ferrotran^®^ and Magtrace^®^ are two iron oxide nanoparticles that are being studied in various cancers. Their diagnostic value in cancers is being established—Magtrace^®^ for the detection of lymph nodes in breast cancer and Ferrotran^®^ in conjunction with MRI. However, the spread of these nanoparticles must be investigated before they can be deemed suitable for MMUS.

In this study, we analyzed the spread of Ferrotran^®^ and Magtrace^®^ to and within lymph nodes in rat after subcutaneous injection to understand their transport, dissemination, and mechanisms as a subcutaneously administered contrast agent. These particles, sized for uptake into the lymphatic system alongside normal draining lymph flow from the injection site, were found in macrophage-like cells and the intercellular space in lymph nodes. Mechanistically, the nanoparticles filled the subcapsular sinus of lymph nodes before entering the inner sinuses. These data suggest that after subcutaneous injection, the nanoparticles transit through the lymph system via the afferent lymphatic vessels into draining lymph nodes. The particles were also found to diffuse into the tissue adjacent to the injection site. Our data confirm the findings of McCauley *et al*. [19] that nanoparticles spread to nearby and distal lymph nodes after subcutaneous administration, as has been observed after intravenous administration, again indicating that these nanoparticles enter the lymphatic system after subcutaneous injection.

With regard to the dynamics of spreading, considering our findings overall, Ferrotran^®^ seems to have dispersed faster than Magtrace^®^, as seen in the macroanalysis of the lymph nodes, which were patently darker after injection of Ferrotran^®^ at equivalent time points. Further, by PPB staining, a higher percentage of cells contained iron after the administration of Ferrotran^®^, particularly after 24 hours, compared with Magtrace^®^. Only Ferrotran^®^ entered the mandibular lymph node after 24 hours, also suggesting more rapid spreading of Ferrotran^®^, as well as the spread of Ferrotran^®^ through the bloodstream. It would have been interesting to determine the time point at which Magtrace^®^ reaches the mandibular lymph nodes; thus, a limitation of this study was that we concluded the experiments at 24 hours and did not measure the spread of iron oxide nanoparticles at later times.

The distance from the injection site to the superficial inguinal nodes in rat is approximately 4 cm versus 12 cm to the axillary lymph node—thus similar to the length of the rectum in human (12–15 cm). Further, the number of lymph nodes that were detected by MRI after subcutaneous administration of Ferrotran^®^ was identical 1 and 3 days after administration, and after 7 days, fewer lymph nodes could be identified [19]. Collectively, these data have implications for imaging, necessitating the optimal dose and time window for administering Ferrotran^®^, which we have attempted to determine in this study.

To be able to detect metastases (or their absence) in lymph nodes, it is desirable to fill most of the lymph node with nanoparticles, rather than only populating the periphery of the lymph node. By PPB staining, at 24 hours post-dose, Ferrotran^®^ filled 10% to 55% of the lymph node area (except for the mandibular lymph node), compared with 2% to 30% with Magtrace^®^, indicating that at least 24 hours is needed to achieve adequate filling. Whether this time is sufficient for MMUS in a clinical setting will need to be tested in a clinical investigation. Our indication from this study, however, is that 24 hours is the minimal required time.

Similarly, biodistribution studies with intravenous administration of Ferrotran^®^ in rats have shown that iron concentrations peak at 24 hours and decrease several days post-dose [25], due to breakdown of the particles to iron by the body’s metabolism of iron and subsequent distribution of iron to the blood and excretion of the dextrose coating in the urine. The optimal time point for imaging with locally administered Ferrotran^®^ is thus expected to be 24–72 hours post-dose—ie, when most of the injected material has entered the lymph nodes but before any significant amount has been metabolized.

We found that subcutaneous Ferrotran^®^ and Magtrace^®^ at the administered dose cause no acute toxicity in rats, consistent with what has been reported for these and other similar nanoparticles [14–17,19], although most studies on Ferrotran^®^ have administered it by intravenous injection. Magtrace^®^ is available on the market, and Ferrotran^®^ has advanced to Phase III trials. Sjins *et al*. [16] found that a dose of Ferrotran^®^ at 2.5 mg Fe/kg body weight (175 mg Fe/70 kg) is safe in human. Also, subcutaneous Magtrace^®^ at a dose of 55 mg Fe has no toxic effects. McCauley *et al*. [19] injected 9 patients subcutaneously in various areas of the pelvis at a dose of 19.6 mg/70 kg in total and observed no severe toxicity.

In a continuation of these toxicity studies, we wanted to analyze the safety of subcutaneous Ferrotran^®^ and Magtrace^®^ in rat with regard to their local and systemic effects. In our study, subcutaneous administration of 11 mg Fe—which is a high local dose for rat (250 g), corresponding to 44 mg/kg body weight—had no local or systemic adverse effects after 24 hours.

This study systematically analyzed the lymph nodes following subcutaneous injection of nanoparticles in a relevant in vivo system, complemented by a detailed characterization of the levels and localization of the tracers in the lymph nodes. This study provides important insights that support the future expansion of these nanoparticles to untested clinical applications.

In humans with rectal cancer, the tracer would be injected under the submucosa near the tumor, but we can not be certain that nanoparticle movement through the tissues in the mesorectum is the same as in subcutaneous tissues, which could affect the time to or amount of uptake by the lymphatic system. However, once the tracer enters the lymphatic system, we would expect their transport via the lymphatic vessels into lymph nodes to be similar; thus, our findings in rats can still be generalized to lymphatic transport in the mesorectum in humans.

This study has several limitations. One drawback was that our study was performed in rats, not humans—between which there are differences with regard to lymph nodes and method of administration (ie, subcutaneous injection in the tail vs the actual clinical diagnostic route) that were not examined in this study. Clinical studies should be performed to determine how Ferrotran^®^ and Magtrace^®^ circulate in the relevant tissues in humans. Nevertheless, studying these tracers in rats has allowed us to dissect lymph nodes at various times after injection, providing valuable insights into their spread and localization in lymph nodes and forming the basis for future clinical studies. We did not conduct an appropriate toxicology study using specific measures of health and behavior, which could be addressed in a subsequent study.

Other limitations of this study include the small sample size. Further, we did not administer Ferrotran^®^ and Magtrace^®^ into animals that bore tumors, which differ from normal tissue in many aspects, potentially influencing the penetration.

Further, we did not compare the lymph node staining and progression of Ferrotran^®^ and Magtrace^®^ against nanocolloids that are used in radionuclide scintigraphy. Finally, our study was primarily qualitative. Future studies that incorporate a quantitative analysis of nanoparticle diffusion and distribution, particularly versus other nanoparticles, will provide additional information. Nevertheless, our findings are sufficient for selection of a candidate tracer for further development and will be confirmed in humans in future studies.

MMUS is a promising diagnostic method for rectal cancer staging, and our results contribute to validating its clinical application. Compared with MRI, MMUS has the potential to be easier to perform and less expensive, uses a smaller device, and requires less training, but most notably, it is potentially more sensitive, as will be shown in a clinical investigation. Once evidence of the value of MMUS for rectal cancer is demonstrated in clinical studies, it could be implemented in other diseases and applications that are monitored using tracers, such as prostate cancer, stem cell therapy, and arteriosclerosis.

## Conclusions

Our study shows that Magtrace^®^ and Ferrotran^®^ reach the proximal lymph nodes at 1 hour post-injection. Over time, their spread within lymph nodes became more homogeneous, and the nanoparticles were observed in distal lymph nodes (axillary), approaching optimal levels at 24 hours post-injection. However, only Ferrotran^®^ reached the mesenteric and mandibular lymph nodes. Thus, we conclude that one should wait at least 24 hours after subcutaneous administration of nanoparticles before imaging the lymph nodes with MMUS. In addition, Ferrotran^®^ spreads faster than Magtrace^®^ throughout the lymphatic system, indicating that Ferrotran^®^ is a potentially more suitable particle for MMUS.

## Abbreviations

ICP-OES: inductively coupled plasma-optical emission spectrometry
MMUS: magnetomotive ultrasound
MRI: magnetic resonance imaging
PPB: Perls Prussian Blue
SPIO: superparamagnetic iron oxide
USPIO: ultrasmall superparamagnetic iron oxide
US: ultrasound

## Ethics approval and informed consent

This study was conducted in accordance with license numbers 18164-21 and M140-16 per the Malmö/Lund Ethics Committee on Animal Testing.

## Consent for publication

Not applicable.

## Data availability

Data will be provided upon reasonable request.

## Authors’ contributions

All authors have read and approved the manuscript. UA analyzed the data, interpreted the results, generated the conclusions, and wrote the manuscript. SB planned and performed the study and analyzed the data. AM planned the study and performed the experiments. LP oversaw the study, planned the study, and wrote the manuscript.

## Acknowledgments

Technical assistance was provided by Micromorph and Timeline Bioresearch. ICP-OES was performed by Spago Nanomedical. Medical writing assistance was provided by Advansci Research Solutions.

